# Dendritic spikes in hippocampal granule cells are necessary for long-term potentiation at the perforant path synapse

**DOI:** 10.1101/253054

**Authors:** Sooyun Kim, Yoonsub Kim, Suk-Ho Lee, Won-Kyung Ho

## Abstract

Long-term potentiation (LTP) of synaptic responses is essential for hippocampal memory function. Perforant-path (PP) synapses on hippocampal granule cells (GCs) contribute to the formation of associative memories, which are considered the cellular correlates of memory engrams. However, the mechanisms of LTP at these synapses are not well understood. Due to sparse firing activity and the voltage attenuation in their dendrites, it remains unclear how associative LTP at distal synapses occurs. Here we show that NMDA receptor-dependent LTP can be induced at PP-GC synapses without backpropagating action potentials (bAPs) in acute rat brain slices. Dendritic recordings reveal substantial attenuation of bAPs as well as local dendritic Na ^+^ ‐spike generation during PP-GC input. Inhibition of Na^+^ ‐spikes impairs LTP suggesting that LTP at PP-GC synapse requires local Na ^+^ ‐spikes. Thus, dendritic spikes are essential for LTP induction at PP-GC synapse and may constitute a key cellular mechanism for memory formation in the dentate gyrus.

## Introduction

The cortico-hippocampal circuit is implicated in the formation, storage, and retrieval of spatial and episodic memories (Lisman, 1999). The dentate gyrus (DG), the first stage in the hippocampal circuitry, receives abundant excitatory projections from the entorhinal cortex via the perforant-path (PP) synapses. Theoretical models of hippocampal function propose that the DG is critically involved in pattern separation and that synaptic transmission and plasticity at PP-granule cell (GC) synapses in the DG is required to remove redundant memory representations (Marr, 1971; McNaughton and Morris, 1987; Treves & Rolls, 1994). In agreement with theoretical predictions, knockout of *N*-methyl-D-aspartate (NMDA) receptors in GCs impairs long-term potentiation (LTP) and the ability to rapidly form a contextual representation and discriminate it from previous similar memories in a contextual fear conditioning task (McHugh et al., 2007). Moreover, GCs that were activated by contextual fear conditioning, referred to as memory engram cells, present clear signatures of synaptic potentiation such as a larger AMPA-NMDA ratio and a greater density of dendritic spines (Ryan et al., 2015). Thus, knowledge of plasticity at PP-GC synapses is essential for understanding the hippocampal function.

Attempts to understand synaptic plasticity rules at PP-GC synapses date to studies of Bliss and Lomo (1973), who first demonstrated LTP in the hippocampus by high-frequency stimulation of the PP fibers *in vivo*. More recently, theta-burst high-frequency stimulation (TBS) of the PP has been used to induce LTP at PP-GC synapses (Schmidt-Hieber et al., 2004; McHugh et al., 2007; Ge et al., 2007). However, the underlying mechanisms for the induction of LTP with TBS at these synapses remain unclear. Associative forms of synaptic plasticity depend on a presynaptic activity (e.g. excitatory postsynaptic potential, EPSP) and a postsynaptic signal (e.g. action potential, AP). Classically, a backpropagating AP (bAP) provides the associative postsynaptic signal at the synaptic site for the induction of LTP (Hebb, 1949; Magee & Johnston, 1997; Dan & Poo, 2006; Feldman, 2012). However, axosomatic APs are poorly propagated back into the dendrites of the GCs (Krueppel et al., 2011), and are unlikely to contribute to the induction of LTP. In addition, mature GCs are relatively silent during exploration (Schmidt-Hieber et al., 2014; Diamantaki et al., 2016) and fire with a low number of APs. Thus, spike-timing dependent plasticity (STDP) that depends on an axosomatic postsynaptic AP will rarely occur under natural conditions (Feldman, 2012). The pronounced attenuation of AP backpropagation, the low occurrence of APs, along with the distinct intrinsic features of mature GCs, such as a hyperpolarized resting membrane potential and reduced excitability (Scharfman & Schwartzkroin, 1990; Mongiat et al., 2009; Pernía-Andrade & Jonas, 2014), cannot explain how distal synaptic PP inputs can be potentiated. Resolving this question has critical implications for understanding both the mechanism and function of memory encoding and retrieval (Ryan et al., 2015).

Understanding LTP induction at PP synapses innervating distal dendrites of GCs requires a detailed analysis of dendritic electrogenesis (Sjöström et al., 2008). Synaptic plasticity at distal synapses may occur in the absence of axosomatic APs via local dendritic spikes (Golding et al., 2002; Kim et al., 2015). In GCs, linear somatic responses can be seen when glutamate is applied in the dendrites (Krueppel et al., 2011). However, without directly accessing the electrical properties of the distal GC dendrites, it is still not known whether synaptic stimulation can generate local dendritic spikes in GCs.

To address this question, we performed subcellular patch-clamp recordings on the thin dendrites of GCs. We found that inhibition of dendritic Na^+^ channels prevented LTP induction via TBS at PP-GC synapses. Because dendritic Na^+^ channels could generate local spikes that were independent of axosomatic AP generation and were not affected by voltage attenuation, we suggest that Na^+^ spikes in the dendrites provide the postsynaptic signal necessary for the induction of LTP at PP-GC synapses.

### Results

We examined GCs with an input resistance (R_in_) < 200 MΩ (R_in_ = 108.4 ± 2.6 MΩ; resting membrane potential: –81.3 ± 0.2 mV; see ‘Materials and Methods’) which corresponds to the mature GC population in the acute hippocampal slices from rats (Schmidt-Hieber et al., 2004).

### TBS-induced LTP at the PP-GC synapses does not require postsynaptic bAPs

We first induced long-term potentiation (LTP) in GCs by theta-burst stimulation (TBS) of the PP synapses in the outer third of the molecular layer (**Figure 1A**). To activate the PP synapses, the tip of a stimulation electrode was placed in close proximity to the dendrite (< 50 μm) under the visual guidance of fluorescent image of the dendrite (‘Materials and Methods’). To ensure that no axosomatic AP initiation and backpropagation occur during TBS, we locally applied tetrodotoxin (TTX) to the GC axon, soma, and proximal dendrites in a subset of experiments (6 out of 13 experiments). TBS-induced LTP caused an increase in the amplitude of the excitatory postsynaptic potentials (EPSPs; **Figure 1B,C**; from 6.96 ± 0.40 mV to 9.96 ± 0.86 mV, n = 13, *P* < 0.01). During TBS, activation of the PP synapses often produced voltage responses with a fast depolarizing phase similar to somatic events observed during dendritic spike generation in other types of neurons (**Figure 1D**; Golding & Spruston, 1998; Golding et al., 2002; Jarsky et al., 2005; Losonczy et al., 2006; Kim et al., 2012). Consistent with previous results (Golding & Spruston, 1998; Golding et al., 2002; Losonczy et al., 2006; Kim et al., 2015), various shapes of putative dendritic spikes were observed during TBS (**Figure 1D**). By examining the temporal derivative (d*V/*d*t*) of somatic voltage responses, we identified weak (d*V*/d*t* < 10 mV/ms) and strong (d*V*/d*t* > 10 mV/ms) putative dendritic spikes that were distinguishable from EPSPs without any local regenerative events (Without dendritic spikes, black, 1.3 ± 0.01 mV/ms, n = 2498; weak dendritic spike, orange, 4.1 ± 0.4 mV/ms, n = 18; strong dendritic spike, red, 14.5 ± 0.8 mV/ms, n = 11; Kruskal-Wallis test: *P* < 0.0001; **Figure 1D**). These putative dendritic spikes were accompanied by a sustained plateau potential (**Figure 1D**; see also **Figure 2B**). Interestingly, the presence of these weak and strong putative dendritic spikes during TBS was correlated with LTP induction (**Figure 1E**; in the presence of putative dendritic spikes, 173.2 ± 12.1%, n = 7; in the absence of putative dendritic spikes 114.1 ± 12.8%, n = 6, *P* < 0.005). Indeed, we found the strong correlation between the number of putative dendritic spikes observed during TBS and the magnitude of LTP (**Figure 1F**; r = 0.77; *P* < 0.005; n = 13). To test the contribution of bAPs in this form of LTP, we applied strong synaptic stimulation without perisomatic TTX application (**Figure 1–figure supplement 1A**). The presence of axosomatic spikes did not significantly affect the magnitude of LTP indicating that bAPs are not critical for this form of LTP (**Figure 1–figure supplement 1B,C**; 167.9 ± 26.8%, n = 8, *P* = 0.87, compared to control in **Figure 1I**). These results suggest that dendritic spikes but not axosomatic APs contribute to LTP induction at PP-GC synapses.

**Figure 1.**
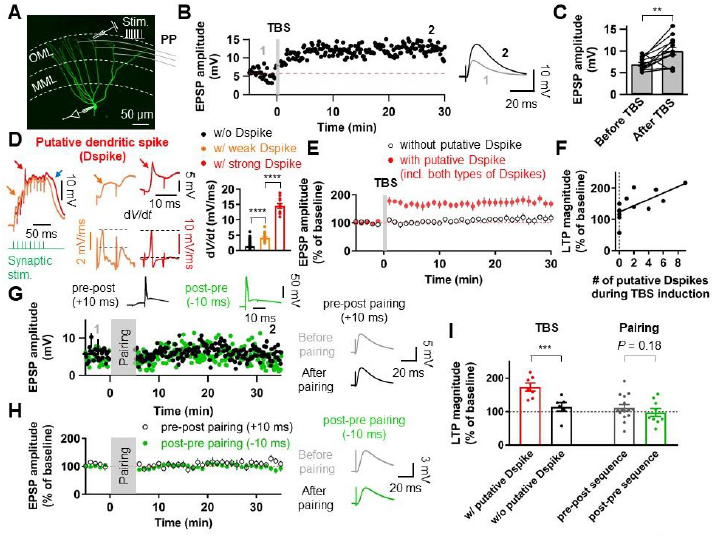
Putative dendritic spikes during theta-burst stimulation (TBS) induction are required for long-term potentiation (LTP) at the perforant-path (PP) to granule cell (GC) synapse. (**A**) Maximum intensity projection of confocal stack fluorescence images of a GC indicating the medial molecular layer (MML) and the outer molecular layer (OML). Synaptic responses of the PP were evoked by electrical stimulation in the OML. Scale bar is 50 μm. (**B**) Time course of excitatory postsynaptic potentials (EPSPs) before (average of 30 EPSP traces, gray) and after (average of 30 EPSPs, black) TBS of the PP synapses. Red line denotes average EPSP baseline value. Increment in EPSP amplitude denotes LTP. (**C**) Bar plot of average EPSP amplitude before and after TBS indicating a significant increment in synaptic responses 25–30 minutes after TBS stimulation (*n* = 13, \*\**P* < 0.01). Data points from the same experiment are connected by lines. (**D**) (Left) Representative somatic voltage traces evoked by high-frequency burst stimulation to the PP synapses. The arrows indicate weak (orange) and strong (red) putative dendritic spikes and accompanying plateau potentials (blue arrow). (Middle) Somatic voltages (top row) and corresponding d*V*/d*t* (bottom row) of weak (orange) and strong (red) putative dendritic spikes in the burst responses (left) on an expaned time scale. Note that d*V/*d*t* values of the putative dendritic spikes increase sharply in a non-linear manner. (Right) Summary bar graphs of d*V/*d*t* peak values of somatically recorded EPSPs with or without dendritic spikes. (**E**) Average time course of EPSPs when TBS stimulation evokes putative dendritic spikes and in the absence of putative dendritic spikes during TBS (black). LTP is induced only if putative dendritic spikes are present during TBS stimulation. (**F**) The number of putative dendritic spikes are significantly correlated with the magnitude of LTP. Black lines represent linear regressions (n = 13). (**G**) Representative time courses of EPSP amplitudes before and after pairing presynaptic stimulation of the PP synapses and postsynaptic action potentials with short time intervals (+10ms, Pre–post sequence, black, top left inset; –10 ms, Post–pre sequence, green, top right inset). Both pairing protocols induce no significant changes in EPSP, suggesting that action potential (AP) backpropagation is not necessary for LTP induction. Right inset shows that LTP was not induced after pairing of EPSPs and APs in both pre-post (top) and postpre sequences (bottom). (**H**) The average EPSP time courses of pre-post (black) and post-pre (green) induction protocols of pairings between synaptic responses and postsynaptic APs, showing that low frequency pairing protocols failed to induce LTP. (**I**) Bar summary graph and individual experiments (circles) indicating that experiments showing the occurrence of putative dendritic spikes during TBS induction induced a significant increment of LTP (i.e., percentage of EPSP baseline), whereas pairings of synaptic stimulation with postsynaptic APs did not show a statistically significant LTP induction, independent of temporal order. Bars indicate mean ± SEM; circles represent data from individual cells. Lines connect data points from the same experiment. \*\*\**P* < 0.005. \*\*\*\**P* < 0.0001. Single-cell data (**B,E**) and mean data (**D,F**; mean ± SEM). Vertical gray bars in **B, D, E,** and **F** indicate the time point of the induction protocol. The following figure supplements are available for figure 1: **Figure supplement 1.** TBS-induced LTP at PP-GC synapses is independent of axosomatic APs. **Figure supplement 2.** Pairing protocols did not induce LTP at the medial PPGC synapses.

To further test whether axosomatic spikes contribute to the induction of LTP at the PP-GC synapse, we used associative pairing protocols. Presynaptic activity was either followed (pre-postsynaptic sequence) or preceded (post-presynaptic sequence) at a 10-ms interval by single or AP bursts (2 APs at 100 Hz). Both protocols failed to induce significant changes in EPSP amplitude (**Figure 1G,H**; pre-postsynaptic sequence, control: 4.97 ± 0.38 mV, after induction: 5.83 ± 0.86 mV, n = 15, *P* = 0.23; post-presynaptic sequence, control: 4.78 ± 0.54 mV, after induction: 4.91 ± 0.99 mV, n = 9, *P* = 0.54), suggesting that APs of axonal origin are not important for potentiation. We further performed similar experiments at the medial PP–GC synapses by stimulating axons in the middle third of the molecular layer (**Figure 1–figure supplement 2A**). Pairing EPSPs and APs in both pre-postsynaptic and post-presynaptic sequences did not induce LTP of EPSPs (**Figure 1–figure supplement 2B–E**; pre-postsynaptic sequence, control: 4.86 ± 0.40 mV, after induction: 5.09 ± 0.99 mV, n = 7, *P* = 0.9015; post-presynaptic sequence, control: 3.86 ± 0.26 mV, after induction: 4.27 ± 0.57 mV, n = 6, *P* = 0.56), consistent with the previous studies that reported no synaptic potentiation after similar pairing protocols (Yang & Dani, 2014; Lopez-Rojas et al., 2016). Together, these results imply that dendritic spikes, rather than axosomatic APs, are the essential signal for LTP induction at distal GC synapses.

### LTP by TBS at PP–GC synapses requires NMDARs and Na ^+^ channels

We next examined the receptors involved in LTP expression at the PP-GC synapses. Bath application of the NMDAR antagonist DL-AP5 (50 μM) abolished LTP (**Figure 2A,D,E**; control: 8.89 ± 0.95 mV; after induction: 9.88 ± 1.21 mV, n = 9, *P* = 0.50). While sustained plateau potentials during LTP induction were inhibited by DL-AP5 in the external solution, fast rising events were remained unchanged (**Figure 2B**). We thus tested whether dendritic voltage-gated Na^+^ channels are involved in LTP. Including 5 mM QX-314 in the whole-cell patch pipette abolished both plateau potentials and TBS-induced LTP (**Figure 2B–D**; control: 11.5 ± 2.5 mV, after induction: 10.6 ± 2.4 mV, n = 8, *P* = 0.38). These results indicate that TBS-induced LTP in GCs depends on the activation of NMDARs and voltage-gated Na^+^ channels on the postsynaptic dendritic membrane, reinforcing the idea that dendritic Na^+^ spikes may play a pivotal role in TBS induction at distal synapses.

**Figure 2.**
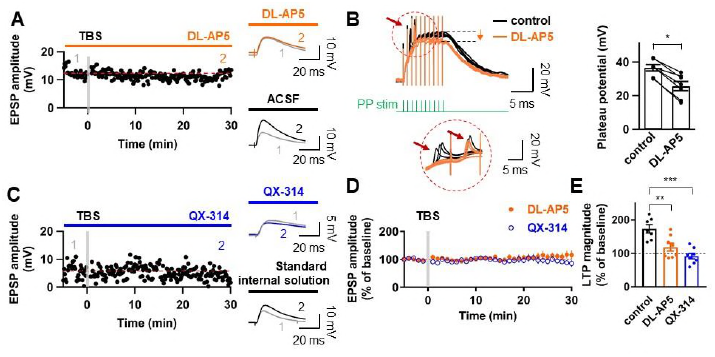
Induction of LTP at PP–GC synapses requires activation of NMDARs and the involvement of Na^+^ channels. (**A**) An example of the time course of EPSP amplitude when TBS protocols was applied in the presence of the NMDAR antagonist, DL-AP5 (50 μM). Red line indicates the average EPSP amplitude. Insets show that EPSP did not increase in the presence of DL-AP5 (orange) after TBS, but a robust LTP was induced when TBS was applied in standard saline (ACSF, black). (**B**) (Left) Long-duration plateau potentials are mediated by NMDA receptor channels. Somatically recorded voltages in response to high-frequency burst sitmulation (green, 10 shocks, 100 Hz) under control (black) and in the presence of DL-AP5 (50 μM; orange). The inset shows putative dendritic spikes (indicated by the red arrow) before and after the addition of DL-AP5; Note that putative dendritic spikes are resistant to the NMDAR blockers. (Right) Summary of the effects of DL-AP5 on plateau potentials. Peak amplitdue of plateau potentials were measured after the stimulus (indicated by dashed lines; Control: 36.6 ± 2.2 mV; DL-AP5: 25.6 ± 3.2 mV; n = 6, \**P* < 0.05). (**C**) A representative time course of EPSP amplitude before and after TBS when the cells were dialyzed with a sodium-channel blocker, QX-314 (5 mM). Inset shows that the averaged EPSP amplitude did not change when blocking sodium channels with QX-314 (blue) despite TBS induction. In contrast, when cells were dialyzed with the standard intracellular solution, the amplitude of EPSP increased after TBS (i.e., LTP). (**D**) Summary plot of TBS-induced LTP experiments in the presence of DL-AP5 (orange) and with dialysis of QX-314 in the recording pipette (blue). Both treatments prevented the induction of LTP at the PP to GC synapse. (**E**) Summary bar graph and individual average EPSP amplitudes after TBS in control (standard saline, black), bath application of DL-AP5 (orange) and dialysis of QX-314 (blue). Treatments with DL-AP5 and QX-314 showed a significant difference compared to the standard LTP induction (DL-AP5, \*\**P* < 0.01; QX-314, \*\*\**P* < 0.005; compared to control in **Figure 1I**). Representative traces in **A** and **C** correspond to the numbers (1 & 2) n the time-course plot. Bars indicate mean ± SEM; circles represent data from individual cells. Lines connect data points from the same experiment. Single-cell data (**A,C**) and mean data (**D**; mean ± SEM). Vertical gray bars in **A, C,** and **D** indicate the time point of the induction protocol.

### Backpropagation of axosomatic APs in the dendrites of GCs

To directly dissect the dendritic mechanism that determines the induction rule of synaptic plasticity in GCs, we employed subcellular patch-clamp techniques to analyze AP backpropagation and initiation in GC dendrites (**Figure 3A**). First, we characterized backpropagation in GCs with somatic current injection evoking trains of APs at the soma while simultaneously recording in the dendrite (**Figure 3B**). The AP always appeared first in the somatic recording and then in the dendrites (**Figure. 3C**). Similar to the previous report (Krueppel et al., 2011), the peak amplitude of the bAPs attenuated as a function of distance from the soma (**Figure 3D,E**). At dendritic distances beyond 150 μm from the soma, the peak amplitude of bAPs was reduced to ∼36% of the amplitude of somatic APs (soma: 93.6 ± 1.7 mV; dendrite: 33.7 ± 3.1 mV; n = 10, *P* < 0.005; **Figure 3D**). The attenuation per dendritic length is much more pronounced in GCs compared to the other neocortical (Stuart et al., 1997; Nevian et al., 2007) and hippocampal pyramidal neurons (Spruston et al., 1995; Kim et al., 2012). It is interesting to note that the extent of bAP attenuation when normalized to the overall length of GC dendrites (mean dendritic length: 278 ± 7.4 μm; n = 11) is quite similar to layer 5 pyramidal neuron dendrites (cf. Figure S3B in Nevian et al., 2007; see also Krueppel et al., 2011; **Figure 3F**), suggesting that the failure of LTP induction by pairing protocol at distal GC synapses (**Figure 1G–I**) might be explained by the voltage attenuation of bAPs, as was reported in layer 5 pyramidal neurons (Letzkus et al., 2006; Sjöström et al., 2008; Feldman; 2012). We also estimated the conduction velocity of bAPs by analyzing the AP latencies measured at the half-maximal amplitude of the rising phase. APs were initiated axonally and propagated back into the dendrites with a velocity of 226 μm/ms (**Figure 3G**; n = 56; corresponding to 0.2–0.3 m/s, Senzai & Buzsaki, 2017). This conduction velocity is slower than those measured in the apical and basal dendrites of layer 5 pyramidal neurons (apical dendrites, 508 μm/ms; basal dendrites, 341 μm/ms; Nevian et al., 2007 and Stuart et al., 1997) and is similar or lower than the velocity estimated in the apical dendrite of other hippocampal principal neurons (Spruston et al., 1995; Kim et al., 2012). In summary, these experiments show that bAPs in GCs propagate into the dendrites with substantial voltage attenuation and moderate conduction velocity.

**Figure 3.**
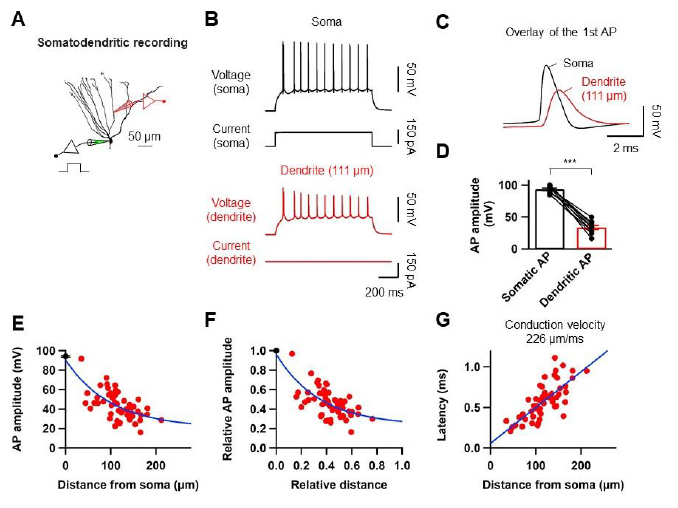
Properties of backpropagating APs in the dendrites of GCs. (**A**) Morphological reconstruction of a GC with representative double somatic and dendritic whole-cell recording configuration used to analyze the AP backpropagation. Scale bar is 50 μm. (**B**) A train of APs elicited by a 1-s current pulse applied at the soma. Black traces indicate somatic voltage and corresponding current; red traces indicate dendritic voltage and corresponding current. (**C**) First AP in the train displayed at expanded time scale. Voltage traces (soma in black, dendrite in red) indicate that the AP is initiated first near the soma and propagated back into the dendrites with a lower amplitude. (**D**) Summary graph to compare somatic (black) and dendritic (red) AP peak amplitude. Bars indicate mean ± SEM; circles represent data from individual cells. Lines connect data points from the same experiment. \*\*\**P* < 0.005 (**E**) Scatter plot of peak amplitude of the backpropagating AP against the absolute physical distance of the recording site from the soma (56 somatodendritic recordings). The blue curve represents a mono-exponential fit to the data points between 0 and 212 μm. (**F**) Scatter plot of the bAP amplitude normalized to the corresponding axosomatic AP amplitude plotted against the distance of the recording site scaled to the total dendritic length (278 ± 7.4 μm; n = 11). The blue curve is a mono-exponential fit to the data. (**G**) Scatter plot of AP latency as a function of the distance from the soma (56 somatodendritic recordings) together with a linear regression (blue line) to compute the average conduction velocity of the AP into the dendrites; dendritic AP propagation velocity was 226 μm/ms. Single-cell data (**E–G**, red) and mean data (**E**, black; mean ± SEM).

### Ionic mechanisms of AP backpropagation

To determine the ionic mechanisms underlying the strong voltage attenuation and the moderate conduction velocity of bAPs, we assayed the somatic and dendritic distribution of voltage-gated Na^+^ and K^+^ conductances in outside-out patches isolated at various locations using pipettes of similar open tip resistance (soma: 17.9 ± 0.5 MΩ, n = 24; dendrite: 19.4 ± 0.5 MΩ, n = 36; **Figure 4**). To calculate conductance density from the measured current amplitude of each component, the surface area of outside-out patches was determined by capacitance measurements (**Figure 4–figure supplement 1**; Schmidt-Hieber & Bischofberger, 2010; Hu & Jonas, 2014). Depolarizing voltage pulses from –120 mV to 0 mV evoked TTX-sensitive inward Na^+^ currents in the majority of outside-out patches excised from both soma and dendrites (**Figure 4A** and **Figure 4–figure supplement 2&3**). Pooled data demonstrated that a moderate density of Na^+^ channels is distributed over the dendritic membrane (**Figure 4D**). On average, the peak Na^+^ conductance density was not significantly different between somatic and dendritic patches (**Figure 4D**, soma, 184.5 ± 36.5 pS/μm^2^, n = 19; proximal dendrite (within 100 μm; PD), 136.4 ± 39.5 pS/μm^2^, n = 8; Distal dendrite (beyond 100 μm; DD), 206.2 ± 38.6 pS/μm^2^, n = 21; soma vs PD and soma vs DD, *P* > 0.99; PD vs DD, *P* = 0.98, Kruskal-Wallis test with Dunn’s multiple comparisons). Surprisingly, a very high density of transient K^+^ currents that appeared to activate and inactivate rapidly in response to voltage pulses from –120 mV to +70 mV was found in patches obtained from the dendrites (**Figure 4B**). These inactivating A-type components were reduced by the A-type K^+^ channel blocker 4-aminopyridine (4-AP, **Figure 4B** and **Figure 4–figure supplement 2&3**). Plotting the conductance density along the dendrite demonstrated that GC dendrites show a significantly higher density of A-type K^+^ components than the soma (**Figure 4E,** soma, 384.8 ± 54.3 pS/μm^2^, n = 21; PD, 784.9 ± 107.9 pS/μm^2^, n = 7; DD, 1319.9 ± 181.6 pS/μm^2^, n = 21; soma vs. PD, *P* = 0.07; soma vs. DD, *P* < 0.0001; PD vs. DD, *P* = 0.76, Kruskal-Wallis test with Dunn’s multiple comparisons). Consistent with this finding, we found that bath application of 5 mM 4-AP in simultaneous somatic and dendritic voltage recordings results in a significant enhancement of the peak amplitude and duration of bAPs in the dendrites, indicating that dendritic A-type K^+^ current (I_A_) contributes to voltage attenuation of bAPs in GCs (**Figure 4–figure supplement 4**). Finally, the delayed rectifier K^+^ current components showed a low and uniform expression over the entire somatodendritic axis (**Figure 4C,F**; soma, 247.9 ± 28.0 pS/μm^2^, n = 21; PD, 231.5 ± 44.3 pS/μm^2^, n = 7; DD, 280.5 ± 46.7 pS/μm^2^, n = 22; *P* > 0.99 for all cases, Kruskal-Wallis test with Dunn’s multiple comparisons). Taken together, these voltage-clamp data reveal that a markedly high density of K^+^ channels and a moderate and uniform density of Na^+^ channels are present in the dendrites of GCs, suggesting the specific biophysical mechanisms underlying both the strong dendritic AP attenuation and the moderate AP propagation velocity.

**Figure 4.**
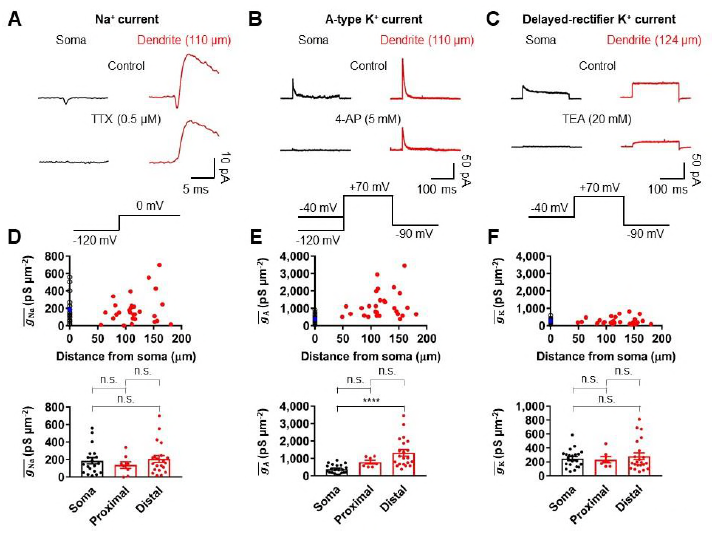
Differential Na^+^ and K^+^ conductance gradients in the dendrites of GCs. (**A**) Averages of Na^+^ current recorded from outside-out patches from soma (black, averages of 25–27 sweeps) and dendrite (red, 110 μm, averages of 20 sweeps) in response to a test pulse potential to 0 mV (bottom). Na^+^ currents were recorded in the presence of 4-AP (5 mM) and TEA (20 mM). Left, soma; right, dendrite; Top, control; bottom, currents in the presence of 0.5 μM TTX in the bath. Leak and capacitive currents were subtracted by a ‘P over –4’ correction procedure. Note that the remaining outward current is the resistant K^+^ current component to 5 mM 4-AP (**Figure 4–figure supplement 2,3**; Hoffman et al., 1997). Blockade of outward K^+^ currents by extracellular 4-AP (5 mM) only had a negligible effects on the peak amplitude of Na^+^ currents (**Figure 4–figure supplement 3**). (**B**) Averages of A-type K^+^ current evoked in outside-out patches excised from soma (black, averages of 6–8 sweeps) and dendrite (red, 110 μm averages of 15–19 sweeps) in response to a test pulse potential to +70 mV (top). Transient A-current was measured by subtraction of traces with a −40 mV prepulse from those with a −120 mV prepulse. Left column, soma; Right column, dendrite; Top row, control; Bottom row, currents in the presence of 5 mM 4-AP in the bath. (**C**) Averages of delayed rectifier K^+^ current evoked in outside-out patches excised from soma (black, averages of 6–8 sweeps) and dendrite (red, 124 μm, averages of 10–18 sweeps) in response to a test pulse potential to +70 mV (top). Delayed rectifier K^+^ current was measured by a −40 mV prepulse. Left column, soma; Right column, dendrite; Top row, control; Bottom row, currents in the presence of 20 mM TEA in the bath. See also **Figure 4–figure supplement 2**. (**D, E, F**) (Top) Plot of Na^+^ channel (**D**), A-type K^+^ channel (**E**), and delayed rectifier K^+^ conductance density (**F**) as a function of distance from the soma, demonstrating that density of various conductances is differentially expressed across the length of GC dendrites. Data from 19, 21, and 21 somatic (black circles) and 29, 28, and 29 dendritic patches (red circles). Blue circles represent the average of somatic recordings. (Bottom) Summary bar graph showing Na^+^ (**D**), A-type K^+^ (**E**) and delayed rectifier K^+^ (**F**) conductance density in the soma, proximal dendrite (< 100 μm) and distal dendrite (≥ 100 μm). Bars indicate mean ± SEM; circles represent data from individual experiments. n.s., not significant; \*\*\*\**P* < 0.0001 by Kruskal Wallis test with *post hoc* multiple comparison using Dunn’s test. The following figure supplements are available for figure 4: **Figure supplement 1.** Estimation of the surface area of outside-out patches by capacitance measurements. **Figure supplement 2.** Pharmacological analysis of voltage-dependent Na^+^ and K^+^ currents. **Figure supplement 3.** The dose-dependent effect of 4-AP on transient outward currents and Na^+^ currents. **Figure supplement 4.** Effect of A-type K^+^ channel blockade on AP backpropagation.

### Dendritically initiated Na ^+^ spikes are required for TBS-induced LTP at PP– GC synapses

Voltage-gated Na^+^ channels may contribute to the generation of dendritically initiated local spikes in regions where the dendrites are very small (Holmes, 1989). Indeed, depolarizing dendrites with brief current pulse injection evoked local spikes in distal dendrites of GCs (55 of 63 recordings; **Figure 5A,B**). In the proximal domain (within 100 μm from the soma), dendritic spikes were not detected and depolarizing dendrites resulted in an axosomatic AP that propagated back to the dendritic recording site (**Figure 5C**). At dendritic recording locations 70 to150 μm from the soma, dendritic spikes were associated with axosomatic spikes. For distances larger than 150 μm from the soma (approximately corresponding to the outer molecular layer), current injections robustly initiated isolated dendritic spikes (**Figure 5C**). Pharmacological analysis revealed that dendritic spikes were resistant to 200 mM CdCl_2_ and 50 mM NiCl_2_ but were inhibited by 0.5 mM TTX, indicating that they were mediated by dendritic voltage-gated Na^+^ channels rather than by Ca^2+^ channels (TTX, 11.3 ± 3.3%, n = 3; CdCl_2_, 93.9 ± 7.8%, n = 4; NiCl_2_, 97.9 ± 10.2%, n = 4; **Figure 5–figure supplement 1**). Because our results indicate that GC dendrites contain a high density of transient, A-type K^+^ channels (**Figure 4B**), we further explored how these channels influence dendritic spike initiation. We combined dual somatodendritic recordings and focal application of 4-AP (10 mM) to the dendritic patch (**Figure 5–figure supplement 2A**). Local block of I_A_ significantly decreased the current threshold for dendritic spike initiation, suggesting that dendritic I_A_ affect the generation of dendritic spikes (**Figure 5–figure supplement 2B,C**).

**Figure 5.**
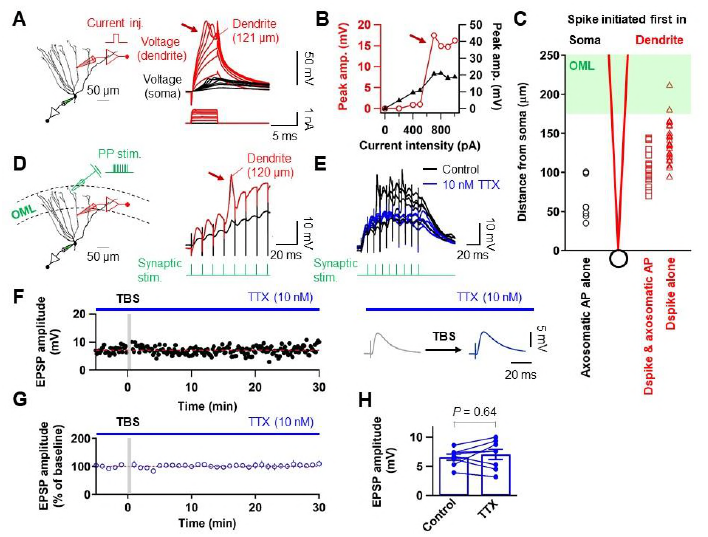
Blockade of Na^+^ channels prevent dendritic spikes and LTP induction by TBS. (**A**) (Left) Schematic recording configuration of a simultaneous somatic (black) and dendritic (red) patch-clamp recording on a GC. (Right) Local dendritic spikes in GCs evoked by dendritic current injection pulses of increasing amplitude (black, voltage in the soma; red, voltage in the dendrite). (**B**) The relationship between voltage amplitude (black, soma; red, dendrite) and dendritic injection resembles a step-function, suggesting the all-or-none nature of the dendritic spike. Peak values of dendritic spikes were measured after subtraction of scaled subthreshold dendritic responses. (**C**) A summary plot showing whether a spike was evoked first in the dendrite (right, red) or in the soma (left, black) for an increasing current pulse injection at the dendrite. Red squares indicate the cells showing a dendritic spike followed by an axosomatic spike. Red triangles show the cells showing isolated dendritic spike. Black circles indicate the cells showing only an axosomatic spike. The green shaded area approximately corresponds to the outer molecular layer. The black circle and red lines in the plot indicate the soma and the dendrite, respectively. (**D**) (Left) Schematic recording configuration of a triple pipette consisting of electrical stimulation of the PP synapses (10 shocks, 100 Hz, green), dendritic patch-clamp recording (120 μm from the soma, red) and somatic whole-cell recording (black). (Right) Dendritic spikes can be identified as larger spikes in the dendrite (red) with the corresponding small spike at the soma (black). (**E**) Bath application of TTX (10 nM) prevents the generation of dendritic spikes evoked by synaptic stimulation (black), as seen on the somatic whole-recording trace (blue). Note that decreasing availability of Na^+^ channels by 10 nM TTX had a negligible effect on the EPSP_1_ (see also **Figure 5–figure supplement 4**). (**F**) Time course of EPSP amplitudes in the presence of TTX (10 nM). The induction of LTP by TBS can be prevented by bath application of TTX, since the average EPSP amplitude does not change. (**G**) Average EPSP before and after TBS shows no changes in amplitude in the presence of TTX. (**H**) Summary bar graph and single experiments (circles) indicating that the TTX application does not produce an increment in the EPSP amplitude and therefore no statistically significant LTP induction. Bars indicate mean ± SEM; circles represent data from individual cells. Data points from the same experiment are connected by lines. Single-cell data (**F**) and mean data (**G**; mean ± SEM). Scale bar in **A** and **D** is 50 μm. Vertical gray bars in **F** and **G** indicate the time point of the induction protocol. The following figure supplements are available for figure 5: **Figure supplement 1.** Dendritic spikes are mediated by voltage-gated Na^+^ channels. **Figure supplement 2.** Dendritic A-type K^+^ channels control dendritic spike initiation. **Figure supplement 3.** Relationship between the d*V*/d*t* of somatically and dendritically recorded voltages during dendritic spikes generation. **Figure supplement 4.** Low concentration of TTX does not affect glutamate release from presynaptic nerve terminals.

We finally examined whether Na^+^ spikes in GC dendrites can be evoked by synaptic stimulation of PP inputs. Extracellular synaptic stimulation was combined with simultaneous double patch-clamp recordings from the soma and dendrites (**Figure 5D**). Dendritic spikes could be triggered by stimulating the PP inputs with TBS protocol (**Figure 5D**) and were inhibited by the application of 10 nM TTX in the bath (**Figure 5E**; Kim et al., 2015). Repeated trials of a high-frequency PP stimulation using the same stimulus intensity produced variable dendritic responses with subthreshold EPSP, weak dendritic spikes, or strong dendritic spikes that appeared either in isolation or associated with axosomatic APs (**Figure 5–figure supplement 3A,B**). When dendritic spikes were present, the d*V*/d*t* of the corresponding somatic voltages showed variable peak amplitudes that were similar to those of the putative dendritic spikes during TBS induction shown in **Figure 1D** (**Figure 5–figure supplement 3C,D**, EPSPs with a dendritic spike: d*V*/d*t* = 7.2 ± 1.1 mV/ms, n = 25; *P* = 0.77 compared to EPSPs with a putative dendritic spike: d*V*/d*t* = 8.0 ± 1.0 mV/ms, n = 29, **Figure 1D**). Finally, low concentration of TTX (10 nM) prevented TBS-induced LTP (**Figure 5E–H**; control: 6.58 ± 0.51 mV; after induction: 7.04 ± 0.88 mV, n = 8; *P* = 0.64), without affecting the baseline EPSP amplitude (**Figure 5–figure supplement 4**; control: 5.21 ± 0.78 mV; after TTX: 5.06 ± 0.68 mV, n = 11; *P* = 0.83). All together, these results suggest that dendritically generated local Na^+^ spikes in response to TBS of the PP are necessary for LTP induction in GCs.

## Discussion

In summary, the present study demonstrates several major findings. First, our results show that conventional STDP protocols (Dan & Poo, 2006; Feldman, 2012) do not trigger synaptic potentiation at PP-GC synapses. Second, a physiologically relevant TBS paradigm can efficiently induce LTP in GCs without axosomatic APs. Finally, direct dendritic recordings revealed that LTP induction requires dendritic spikes. To our knowledge, these studies show the first direct evidence for dendritic spike generation in GCs and the role of these spikes during induction of LTP in the dentate gyrus network. However, whether our findings also hold for GCs at early stages of development remains to be determined.

Considerable evidence has shown that the electrical properties of neurons at the cellular and subcellular levels are brain region-specific and cell type-specific and endow unique rules and characteristics on circuit function (Sjöström et al., 2008; Stuart & Spruston, 2015). Our findings support this idea. Hyperpolarized resting potential, strong dendritic voltage attenuation, and low occurrence of APs are the known electrophysiological characteristics of mature GCs (Scharfman & Schwartzkroin, 1990; Schmidt-Hieber et al., 2004; Mongiat et al., 2009; Krueppel et al., 2011; Pernía-Andrade & Jonas, 2014), which are distinct from other types of hippocampal principal neurons (Spruston et al., 1995; Kim et al., 2012; Spruston & Stuart, 2015). While backpropagated APs are an important associative signal for triggering plasticity at the synaptic site via the voltage-dependent relief of Mg^2+^ block of NMDARs (Magee & Johnston, 1997; Dan & Poo, 2006; Feldman, 2012; Mishra et al., 2016), the above features of mature GCs are highly unfavorable for LTP induction at distal synaptic contacts. Accordingly, we found that GCs do not show LTP at PP-GC synapses during standard STDP (**Figure 1G,H** and **Figure 1–figure supplement 2**), which is consistent with recent reports (Yang & Dani, 2014; Lopez-Rojas et al., 2016).

However, several studies have demonstrated EPSP-AP pairing protocol-induced synaptic potentiation at the same synapses (Levy & Steward, 1983; Lin et al., 2006). Given that Levy and Steward (1983) and Lin et al. (2006) employed *in vivo* and *in vitro* field recording configurations, respectively, the discrepant results could be attributed to GCs at different stages of maturation as immature GCs exhibit a lower threshold for LTP induction (Schmidt-Hieber et al., 2004; Ge et al., 2007) or under different recording circumstances that were exposed to various neuromodulators. For example, Yang and Dani (2014) reported that the pairing protocol that showed no synaptic potentiation could induce reliable LTP at PP-GC synapses after D1-type dopamine receptor activation by which dendritic A-type K^+^ currents are suppressed. Because I_A_ is known to limit the backpropagation of APs (Hoffman et al., 1997), suppresion of I_A_ can boost AP backpropagation, allowing sufficient dendritic depolarization for LTP induction at distal synapses. Our observations of a high density of dendritic I_A_ and their effects on AP backpropagation in GCs directly support the finding of Yang and Dani (2014). Therefore, under physiological conditions, long-lasting dendritic depolarization during theta oscillation (Buzsaki, 2002) or activation of neuromodulatory systems (Hamilton et al., 2010; Yang & Dani, 2014) may cause inactivation of dendritic I_A_ and trigger pairing-induced LTP (Lin et al., 2006; Brunner & Szabadics, 2016).

Although the presence of A-type K^+^ channels in GC dendrites had been shown in earlier immunocytochemical studies (Birnbaum et al., 2004; Monaghan et al., 2008; Menegola et al., 2008), it has been proposed that dendritic A-type K^+^ channels have no significant impact on bAP-induced Ca^2+^ signals (Krueppel et al., 2011). Krueppel et al.’s (2011) results appear to be inconsistent with our present data, which show the strong expression of functional A-type K^+^ channels in GC dendrites. In contrast to that study, we directly tested the effect of 4-AP in simultaneous soma-dendrite recordings by using both global and local application methods. Our results show that the effect of local puff application of 4-AP to the dendritic recording site is much smaller than that of global application. As Krueppel et al. (2011) used local application halfway between the soma and the Ca^2+^ imaging site, the impact of 4-AP on bAPs in their study could be underestimated.

We further found that high-frequency stimulation of PP synapses is efficient in eliciting dendritic spikes required for LTP induction. As reported in a recent *in vivo* study, mature GCs are exposed to abundant functional glutamatergic inputs from the entorhinal cortex during theta rhythm (Pernía-Andrade & Jonas, 2014; Schmidt-Hieber et al., 2014). Moreover, several lines of evidence demonstrated that mature GCs are highly innervated by PP synaptic connections while receiving a powerful perisomatic inhibition (Dieni et al., 2013, 2016; Temprana et al., 2015). Under these *in vivo* network conditions, strong dendritic excitation by high-frequency PP inputs and strong perisomatic inhibition may promote amplification of dendritic responses without axosomatic APs. Therefore, dendritic spikes are likely physiologically relevant signals for the induction of LTP. Support for the idea of cooperative LTP at PP-GC synapses has also come from McNaughton et al. (1978), who demonstrated that high-frequency stimulation of PP synapses could induce synaptic enhancement in the absence of GC discharges.

This specific high-frequency stimulation-dependent synaptic potentiation in GCs presumably stems from distinct functional and geometrical features of GC dendrites. A moderate density of dendritic Na^+^ channels together with higher input impedance of the distal dendrites (Hama et al., 1989; Schmidt-Hieber et al., 2007; Holmes, 1989) suffices for initiating spikes locally in distal dendrites that have small capacitive load but not for supporting active backpropagation of axosomatic APs. Although GCs have relatively short dendrites, a high K^+^ to Na^+^ current ratio in these thin-caliber dendrites imposes a strong distance-dependent attenuation of axosomatic APs, leading to a lack of pairing-induced LTP at distal synapses. Thus, Na^+^ spikes in the dendrites contribute the postsynaptic depolarization necessary for the induction of associative plasticity (Golding et al., 2002; Kim et al., 2015). However, it should be noted that dendritic Na^+^ spikes were accompanied by NMDAR-mediated plateau potentials and therefore these two dendritic events could act in concert to trigger LTP in GCs (Schiller et al., 2000).

Consequently, our findings suggest that in the absence of axonal firing (Alme et al., 2010; Diamantaki et al., 2016), dendritic spikes would allow a silent GC to participate in the storage of memories via LTP induction. It would further permit a functional separation between a storage phase mediated by Na^+^ channels in GC dendrites, and a recall phase (O´Neil et al., 2008) that effectively activates the CA3 neurons (Vyleta et al., 2016) via Na^+^ channels in GC axons.

## Materials and Methods

### Slice preparation and electrophysiology

Acute hippocampal slices (thickness, 350 μm) were prepared from the brains of 17‐ to 25-day-old Sprague-Dawley rats of either sex. Rats were anesthetized (isofluorane, Forane®; Abbott) and decapitated rapidly. All the experiments were approved by the University Committee Animal Resource in Seoul National University (Approval #: SNU-090115-7). All brains were sliced coronally, and dorsal slices displaying all the subregions of the hippocampal formation were used for the experiments (Coronal sections located between 4.3 mm and 5.7 mm from the posterior end of the right hemisphere). Slices were prepared in an oxygenated ice-cold sucrose-containing physiological saline using a vibratome (VT1200, Leica), incubated at ∼36°C for 30 min, and subsequently maintained in the same solution at room temperature until the recordings. Recordings were performed at near-physiological temperature (33–35°C) in an oxygenated artificial cerebral spinal fluid (ACSF).

Patch pipettes were made from borosilicate glass capillaries (outer diameter = 1.5 mm, inner diameter = 1.05 mm) with a horizontal pipette puller (P-97, Sutter Instruments). The open-tip resistance of patch pipettes was 2.5– 6.5 MΩ and 11–30 MΩ for somatic and dendritic recordings, respectively. Current-clamp recordings were performed with an EPC-10 USB Double amplifier (HEKA Elektronik). In current-clamp recordings, series resistance was 8–80 MΩ. Pulse protocols were generated, and signals were low-pass filtered at 3 or 10 kHz (Bessel) and digitized (sampling rate: 20–50 kHz) and stored using Patchmaster software running on a PC under Window 10. Resting membrane potential (RMP) was noted immediately after rupture of the patch membrane. R_in_ was determined by applying Ohm’s law to the steady-state voltage difference resulting from a current step (±50 pA). Pipette capacitance and series resistance compensation (bridge balance) were used throughout current-clamp recordings. Bridge balance was checked continuously and corrected as required. Experiments were discarded if the resting membrane potential depolarized above –70 mV and were stopped if the resting membrane potential or R_in_ changed by more than 20% during the recording.

All experiments were performed on visually identified mature GCs on the basis of the relatively large and round-shaped somata, and the location of the cell body under DIC optics. GCs located at the superficial side of the GC layer in the suprapyramidal blade were preferentially targeted. These cells had the average RMP of –81.3 ± 0.2 mV and R_in_ of 108.4 ± 2.6 MΩ (n = 165), that is similar to characteristic intrinsic properties of mature GC population (Schmidt-Hieber et al., 2004). Cells were filled with a fluorescent dye, Alexa Fluor 488 (50 μM, Invitrogen) and imaged with an epifluorescence system mounted on an upright microscope equipped with a 60 x (1.1 N.A.) water immersion objective lens. Focal electrical stimulation (100 μs pulses of 1–35 V) was applied on isolated dendrites in the outer third of the molecular layer (within 100 μm of the hippocampal fissure) by placing a glass microelectrode (0.5–3 MΩ) containing 1 M NaCl or ACSF in the vicinity of the selected dendrite (typically at < 50 μm distance), guided by the fluorescent image of the dendrite. All experiments were performed in the presence of the GABA receptor antagonist picrotoxin (PTX, 100 μM) and CGP52432 (1 μM).

To record voltage-gated Na^+^ or K^+^ currents, outside-out patches were excised from the soma and the dendrite with pipettes of similar geometry and open-tip resistances (18.8 ± 0.4 MΩ, n = 60, ranging from 12.5 to 24.4 MΩ) for comparison of channel density between soma and dendrite (Kim et al., 2012). Ensemble K^+^ currents were evoked by a pulse protocol consisting of a 50–200 ms prepulse to –120 mV followed by a 200 ms test pulse to 70 mV. Na^+^ currents were generated by a pulse sequence comprised of a 100-ms prepulse to –120 mV and a 30-ms test pulse to 0 mV. In all cases, the holding potential was –90 mV before and after the pulse protocol. Voltage protocols were applied to outside-out patches once every 3 and 5 s for Na^+^ and K^+^ current recordings, respectively. Leak and capacitive currents were subtracted online using the pipette capacitance compensation circuit of the amplifier and a ‘P over –4’ correction procedure (Schmidt-Hieber & Bischofberger, 2010).

### Subcellular dendritic patch-clamp recording

Dendritic recordings from GCs were obtained similarly as described previously (Kim et al., 2012). First, Alexa Fluor 488 (50 μM, Invitrogen) diluted in an internal solution was loaded into cells via a somatic recording pipette. Second, after ∼10 min of loading, fluorescently labeled dendrites were traced from the soma into the molecular layer using epifluorescence microscope. Finally, fluorescent and infrared differential interference contrast (IR-DIC) images were compared, and GC dendrites were patched under IR-DIC.

### Stimulation protocols for the induction of long-term potentiation (LTP)

LTP was induced by either theta burst stimulation (TBS; Schmidt-Hieber et al., 2004) or pairing protocols. TBS induction protocol consisted of burst of EPSPs (10 stimuli at 100 Hz) repeated 10 times at 5 Hz. These episodes were repeated four times every 10 seconds. The pairing protocol comprised 300 repetitions of a presynaptic stimulation and one or two postsynaptic APs at the different time intervals at 1 Hz. A postsynaptic AP was evoked by a brief current injection to the soma (2 ms, 3 nA). For LTP experiments, baseline EPSPs evoked by stimulating presynaptic axon fibers at 0.1 Hz were measured for ∼10 min after whole-cell recording. After LTP induction protocol, EPSPs were recorded for 30 min. LTP magnitude was evaluated as the percentage of EPSP baseline (5 min) after 25 to 30 minutes after the induction protocol. In a subset of recordings, local application of TTX (1 μM) to the perisomatic area was used to prevent axosomatic AP initiation and backpropagation during TBS induction, without affecting evoked EPSPs.

### Solutions and chemicals

The extracellular solution for dissection and storage of brain slices was sucrose-based solution (87 mM NaCl, 25 mM NaHCO_3_, 2.5 mM KCl, 1.25 mM NaH_2_PO_4_, 7 mM MgCl_2_, 0.5 mM CaCl_2_, 10 or 25 mM glucose, and 75 sucrose). Physiological saline for experiments was standard ACSF (125 mM NaCl, 25 mM NaHCO_3_, 2.5 mM KCl, 1.25 mM NaH_2_PO_4_, 1 mM MgCl_2_, 2 mM CaCl_2_, and 25 mM glucose). TTX and 4-AP were applied either via bath perfusion (0.5 μM and 10 mM, respectively) or by local application (1 μM and 10 mM, respectively) with a pressure application system (Picospritzer 3, General Valve). Pressure pulses had durations of 0.2 s and amplitudes of ∼10 psi. CdCl_2_ and NiCl_2_ were applied in the bath at a concentration of 200 μM and 50 μM, respectively.

For whole-cell recording and K^+^ current recording in outside out patches, we used K^+^ rich intracellular solution that contained 115 mM K-gluconate, 20 mM KCl, 10 mM HEPES, 0.1 mM EGTA, 2 or 4 mM MgATP, 10 mM Na_2-_phosphocreatine, and 0.3 mM NaGTP, pH adjusted to 7.2–3 with KOH (∼300 mOsm). In a subset of experiments, 50 μM Alexa 488 and 0.1–0.2 % biocytin (wt/vol) were added to the internal solution for labeling during the experiment or after fixation, respectively. All drugs were dissolved in physiological saline immediately before the experiment and perfused on slices at a rate 4–5 ml min^-1^. These included: DL-AP5, PTX, CGP52432, TTX, 4-AP, and TEA. Intracellularly delivered QX-314 (N-(2,6-dimethylphenylcarbamoylmethyl) triethylammonium chloride) were directly added to the pipette solution before the experiment was started. TTX and DL-AP5 (D,L-2-amino-5-phoshonovaleric acid; 50 μM) were purchased from Tocris Bioscience; CGP52432 were from Abcam; All other drugs were from Sigma-Aldrich.

### Immunohistochemistry

Post hoc morphological analysis of GCs was performed as described previously with slight modifications (Kim et al., 2012). Slices were fixed overnight at 4°C in 4% paraformaldehyde in 100 mM phosphate buffer solution (PBS), pH7.4. After fixation, slices were rinsed several times with PBS and permeabilized with 0.3% Triton X-100 in PBS. Subsequently, slices were treated with 0.3% Triton X-100 and 0.5% BSA in PBS to prevent any unwanted. Next, slices were treated with 0.3% Triton X-100 and 0.2% streptavidin-cy3 in PBS and were again incubated overnight in 4°C. Finally, slices were mounted with DAKO S3023 medium.

## Data analysis

Custom-made routines written in Igor Pro 6.3 (Wavemetrics), Stimfit 0.15 (Guzman et al., 2014) or Prism (Graphpad) were used for data analysis and statistical testing. Spike threshold value was determined as the time point at which the derivative of voltage exceeded 40 V/s at the soma or 20 V/s at the dendrite. To determine the conduction velocity, latency–distance data were fit with a linear regression, and the velocity was calculated from the linear regression slope. Conductance density (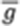) was calculated as 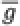 *= I_Peak_ / ((V– V_rev_) • A)*, where *I_Peak_* is the corresponding peak current amplitude, *V* is the test pulse potential, and *A* is the membrane surface area of outside-out patches. Membrane patch area was estimated by capacitance measurement (**Figure 4– figure supplement 1**; Schmidt-Hieber & Bischofberger, 2010; Hu & Jonas, 2014). Reversal potentials were assumed as +75 mV for Na^+^ current and –95 mV for K^+^ currents (Kim et al., 2012).

All values indicate mean ± standard error of the mean (SEM), with “*n*” denoting the number of experiments. To test statistical significance, a two-sided nonparametric Wilcoxon signed-rank test or Wilcoxon rank sum test were used (unless noted otherwise). For comparisons of more than two groups, a Kruskal-Wallis test with Dunn’s multiple comparisons for post-hoc testing was used. Differences with P-value less than 0.05 were indicated with an asterisk and considered significant. Membrane potential values were displayed without correction for liquid junction potentials. Distances were measured from the soma to the dendritic recording site along the dendritic trajectory (Kim, 2014).

## Acknowledgments

We thank Jose Guzman and Hua Hu for critically reading the manuscript.

## Figure supplement

**Figure 1–figure supplement 1.**
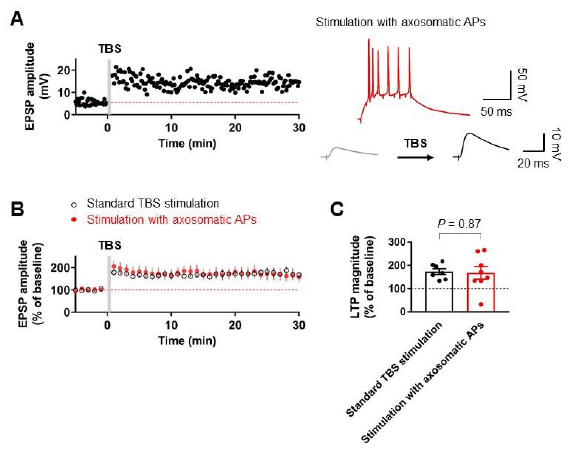
TBS-induced LTP at PP-GC synapses is independent of axosomatic action potentials (APs). (**A**) Time course of excitatory postsynaptic potentials (EPSPs) before (average of 30 EPSP traces, gray) and after (average of 30 EPSPs, black) theta-burst stimulation (TBS) of the PP synapses. Red line denotes average EPSP baseline value. Inset: example of the first burst of TBS responses (red) showing initiation of multiple axosomatic APs during TBS induction upon increasing the stimulus intensity. (**B**) Average time course of EPSPs when TBS stimulation evokes axosomatic APs (red) and in the absence of axosomatic APs during TBS (black). Note that a similar degree of potentiation in both cases is shown. (**C**) Summary bar plot of average EPSP amplitude before and after TBS indicating that there are no significant differences in the magnitude of LTP between two groups in **B**. Bars indicate mean ± SEM; circles represent data from individual cells. Single-cell data (**A**) and mean data (**B**; mean ± SEM). Vertical bars (gray) in **A** and **B** indicate the time point of the pairing protocol.

**Figure 1–figure supplement 2.**
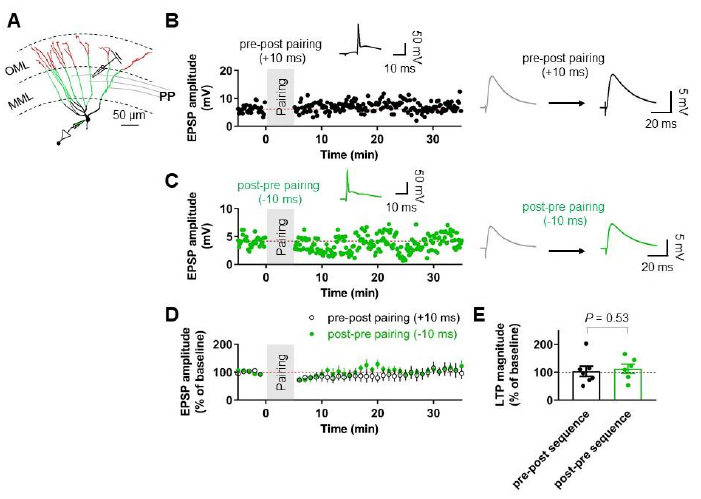
Pairing protocols did not induce LTP at the medial perforant path (MPP)-GC synapses. (**A**) An example of reconstructed granule cells and diagram illustrating the experimental configuration. Note that a stimulating electrode was placed in the middle third of the molecular layer. Scale bar is 50 μm. (**B**) Representative time course of EPSP amplitude, showing that repeated pairing of EPSPs and following postsynaptic APs (pre-post sequence; *Δt* = +10 ms, inset) induced no significant LTP at MPP-GC synapses. (Right) EPSPs before (gray) and 25–30 min after induction (black). (**C**) Representative time course of EPSP amplitude, showing that postpresynaptic pairing (*Δt* = –10 ms, inset) induces no LTP at MPP-GC synapses. (Right) EPSPs before (gray) and 25–30 min after induction (green). (**D**) Summary graph of pairing-induced LTP experiments at the MPP-GC synapses demonstrating the absence of LTP. (**E**) Summary graph and scatter (individual cells) plot of the change in EPSP amplitude before and after the pairing protocols. Bars indicate mean ± SEM; circles represent data from individual cells. Single-cell data (**B,C**) and mean data (**D**; mean ± SEM). Vertical gray bars in **B**, **C**, and **D** indicate the time point of the induction protocol.

**Figure 4–figure supplement 1.**
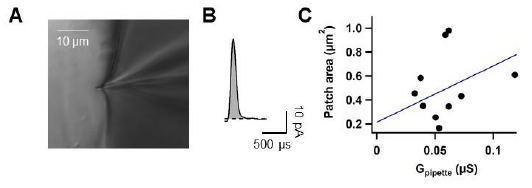
Estimation of the surface area of outside-out patches by capacitance measurements. (**A**) Experimental configuration showing the patch pipette near the silicone elastomer ball (Sylgard 184, Dow Corning). After obtaining excised outside-out patches, the pipette tip was gently pressed against the silicone ball (< 5 μm), resulting in complete sealing of the tip. All the pipettes were coated with dental wax for this set of experiments (Kimsdent). (**B**) The traces show the difference current (evoked by a test pulse from –60 to –110 mV, average of 100 individual capacitive current traces) before and after pressing the pipette tip into the silicone ball. The integral under the curve (grey area) represents the capacitive charge used for membrane surface (Schmidt-Hieber & Bischofberger, 2010). Membrane surface area was determined from the capacitance assuming a specific membrane capacitance of 1.0 μF•cm^-2^. Dashed lines indicate leakage current. (**C**) Plot of membrane surface area of outside-out patches against pipette open-tip conductance. Data (n = 10) were fit by linear regression according to A(g_P_) = 4.7124 x g_P_ + 0.21421, where A is membrane patch area (μm^2^) and g_P_ is pipette conductance (μS).

**Figure 4–figure supplement 2.**
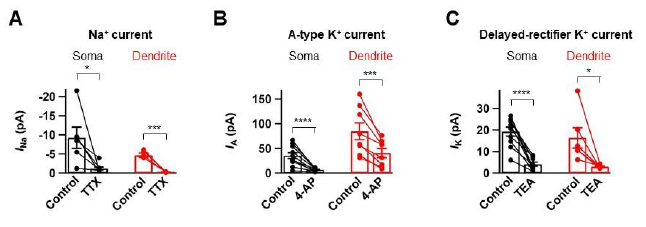
Pharmacological analysis of voltage-dependent Na^+^ and K^+^ currents. (**A**) Bar graph summarizing the effects of the I_Na_ channel blocker TTX (0.5 μM) on the transient Na^+^ currents excised from the soma (left, black, n = 6) and the dendrite (right, red, n = 6). (**B**) Bar graph summarizing the effects of the I_A_ channel blocker 4-AP (5 mM) on the transient outward K^+^ currents excised from the soma (left, black, n = 10) and the dendrite (right, red, n = 8). (**C**) Bar graph summarizing the effects of the I_K_ channel blocker TEA (20 mM) on the steady-state outward K^+^ currents excised from the soma (left, black, n = 10) and the dendrite (right, red, n = 6). *0.01 ≤ *P* < 0.05; **p < 0.01; \*\*\**P* < 0.005; \*\*\*\**P* < 0.0001 by paired t-test. Bars indicate mean ± SEM; circles represent data from individual cells. Lines connect data points from the same experiment.

**Figure 4–figure supplement 3.**
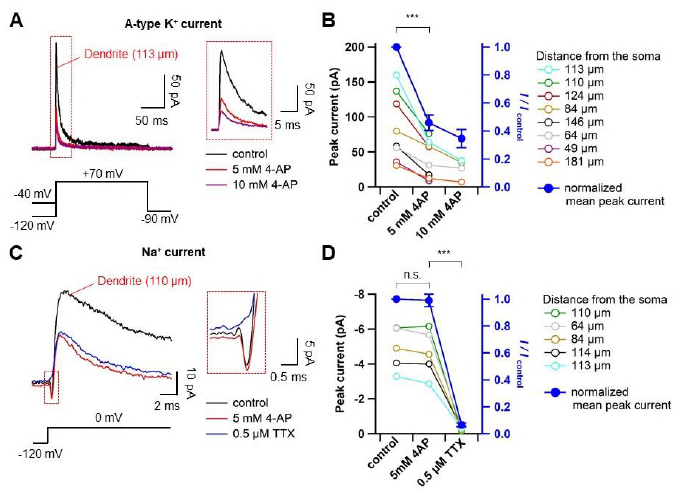
The dose-dependent effect of 4-AP on transient outward currents and Na^+^ currents. (**A**) Representative traces showing dendritic A-type K^+^ currents (113 μm from the soma, averages of 10 sweeps) evoked as in **Figure 4B** in control (black) and subsequently in 5 mM 4-AP (red), followed by 10 mM 4-AP (purple) in the bath. Inset shows an expanded traces corresponding to the dashed box in the left panel. (**B**) Summary plot indicating that 4-AP reduced the transient K^+^ current in a dose-dependent manner. Colored circles represent data from individual dendritic recordings at different distances (right). Lines connect data points from the same experiment. Filled circles (blue) represent average data of the normalized peak currents, showing the percentage block of transient outward K^+^ currents in the presence of 5 and 10 mM 4-AP (right axis; 5 mM 4-AP, n = 8, \*\*\**P* < 0.005 by paired t-test). (**C**) Representative traces showing averages of Na^+^ currents (top, 110 μm from the soma, averages of 10 sweeps) evoked as in **Figure 4A** in control (black) and subsequently in 5 mM 4-AP (red), followed by 0.5 μM TTX and 5 mM 4-AP (blue) in the bath. Inset shows an expanded traces corresponding to the dashed box on the left. (**D**) Summary graph showing the effects of 5 mM 4-AP and 0.5 μM TTX on Na^+^ currents. Colors as in **B**. Filled blue circles show the average of the normalized peak currents demonstrating that 5 mM 4-AP did not significantly affect the peak of transient inward currents (n = 6, *P* = 0.69; n.s., not significant, by paired t-test). These inward currents were blocked by 0.5 μM extracellular TTX (n = 5, \*\*\**P* < 0.005 by paired t-test). Open circles represent data from individual cells and filled circles represent mean data (mean ± SEM).

**Figure 4–figure supplement 4.**
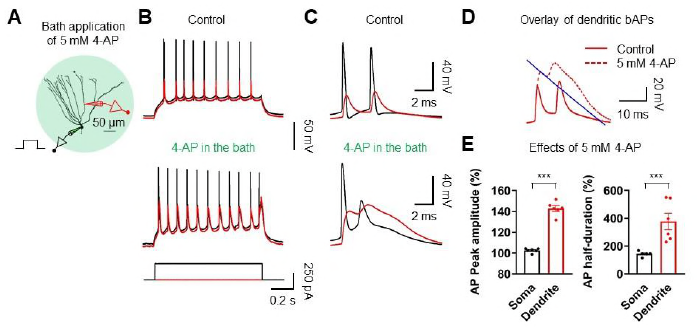
Effect of 4-AP on AP backpropagation. (**A**) Diagram of the experimental setup displaying simultaneous recordings from the soma (black) and the dendrite (red) of a GC in the presence of 5 mM 4-AP. Scale bar is 50 μm. (**B**) Sample traces of somatic (black) and dendritic (red) APs evoked by somatic current injection in the absence (top; control) or the presence of 4-AP (5 mM; bottom) in the bath. (**C, D**) First and second APs (**C**) in the train displayed at expanded time scale and overlay of dendritic voltages (**D**) under control and in the presence of 4-AP (dashed line). The blue line illustrates an exponential fit to the repolarizing phase of the 1st dendritic AP to calculate the duration at half-maximal amplitude. (**E**) Summary graph of the effects of 5 mM 4-AP on AP peak amplitude (Left; Soma: 102.6 ± 0.8%; Dendrite: 142.8 ± 2.6 %; n = 6, ***P < 0.005) and duration at half-maximal amplitude (right; soma: 144.4 ± 7.1%; dendrite: 377.6 ± 59.0%; n = 6, ***P < 0.005) in 6 somatodendritic recordings. Bars indicate mean ± SEM; circles represent data from individual cells.

**Figure 5–figure supplement 1.**
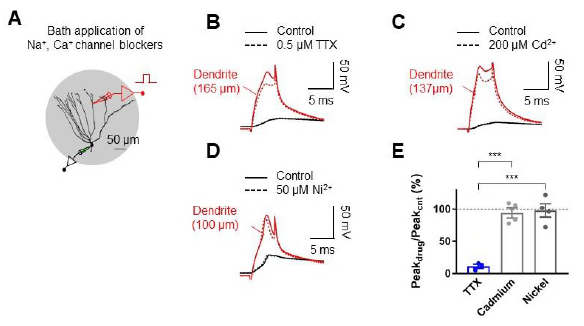
Dendritic spikes are mediated by voltage-gated Na^+^ channels. (**A**) Schematic diagram illustrating the recording configuration of a simultaneous somatic (black amplifier) and dendritic (red amplifier) patch-clamp recording on a GC combined with bath application of blockers of voltage-gated Na^+^ and Ca^2+^ channel. Scale bar is 50 μm. (**B**) Blockade of voltage-gated Na^+^ channels by a bath application of TTX (0.5 μM) eliminated dendritic spikes elicited by a 5-ms current pulse injected into the dendritic electrode. (**C**) Application of 200 μM CdCl_2_ did not abolish dendritic spikes. Bath application of CdCl_2_ did not affect the late depolarizing phase of the dendritic spikes (S.K., unpublished observations). (**D**) Application of 50 μM NiCl_2_ did not abolish dendritic spikes. (**E**) Summary of the effects of the indicated pharmacological agents on the peak of dendritic spikes. The dendritic recording sites are 165 μm (**B**), 137 μm (**C**), and 100 μm (**C**) from the soma, respectively. Bars indicate mean ± SEM; ***P < 0.005 by Student’s ttest; Black traces: somatic voltage; red traces: dendritic voltage.

**Figure 5–figure supplement 2.**
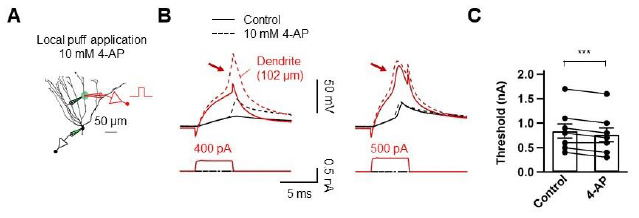
Dendritic A-type K^+^ channels control dendritic spike initiation. (**A**) Diagram of the experimental setup illustrating the recording configuration of a simultaneous somatic (black amplifier) and dendritic (red amplifier) patch-clamp recording on a GC combined with focal application of 10 mM 4-AP directly to the dendritic patch. Scale bar is 50 μm. (**B**) Somatic and dendritic voltage responses to dendritic current injection pulses with increasing amplitude (left: 400 pA; right; 500 pA). Local application of 4-AP to the dendrite near the recording pipette tip decreased the threshold for initiating dendritic spikes (arrows). Black traces represent somatic signal and red traces represent dendritic signal. Dashed line indicates the voltage response after puff application of 4-AP. The dendritic recording site is 102 μm from the soma. (**C**) Summary of the effects of local application of 4-AP on the threshold required to initiate dendritic spikes (Control, 0.84 ± 0.15 nA; 4-AP, 0.76 ± 0.14 nA; n = 8, \*\*\**P* < 0.005, Paired t-test). Bars indicate mean ± SEM. Lines connect data points from the same experiment.

**Figure 5–figure supplement 3.**
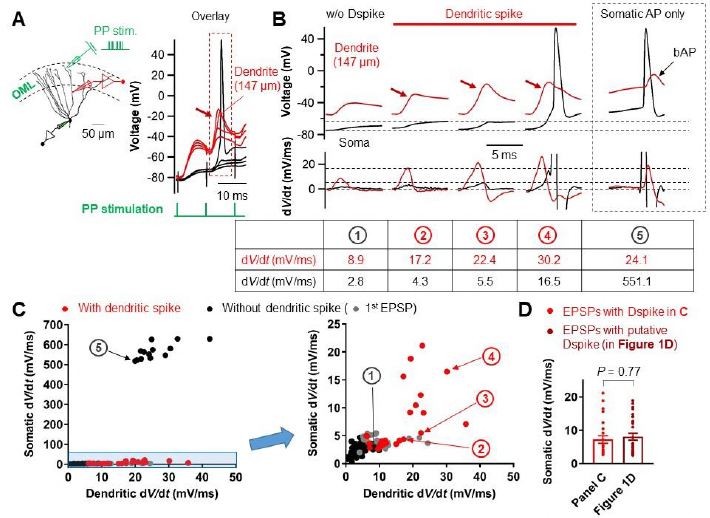
Relationship between the d*V*/d*t* of somatically and dendritically recorded voltages during dendritic spikes generation. (**A**) (Left) Diagram illustrating the experimental configuration of a triple pipette consisting of electrical stimulation of the PP synapses in the OML (green), dendritic patch-clamp recording (147 μm from the soma, red) and somatic whole-cell recording (black). Scale bar is 50 μm. (Right) Somatic and dendritic voltages in response to a high-frequency PP stimulation with a constant stimulus intensity are shown superimposed for comparison. (**B**) Traces of somatic (black) and dendritic (red) voltage responses (top row) in **A** (red box) and corresponding d*V*/d*t* (Bottom). d*V*/d*t* peak amplitudes of each traces were summarized in the table (bottom). The red arrows indicate the dendritic spikes. When dendritic spikes were present, the corresponding somatic voltage changes were used for analysis of d*V*/d*t* peaks. For comparison, somatic and dendritic membrane voltages and corresponding d*V*/d*t* traces during axosomatic AP generation are also shown on the right (dashed box). Encircled numbers indicate correspondence between traces in **B** and data points in **C**. (**C**) (Left) Summary plot of peaks in somatic d*V*/d*t* against corresponding peaks in dendritic d*V*/d*t* for 4 simultaneous somatodendritic recordings in response to high-frequency burst stimulation of the PP synapses (Dendritic recording sites are from 136 μm to 175 μm from the soma). (Right) An enlarged view of the box (blue) in the left panel. Black circles, in the absence of dendritic spikes; red circles, in the presence of the dendritic spikes; gray circles, data points from the first EPSPs (EPSP_1_). Note that somatic d*V*/d*t* peak amplitudes of subthreshold EPSP1s were comparable to those of the somatic traces when dendritic spikes were present. (**D**) Summary bar graphs of d*V/*d*t* peak amplitudes of somatically recorded dendritic spikes in **C** (red) and in **Figure 1D**. Bars indicate mean ± SEM; circles represent data from individual cells.

**Figure 5–figure supplement 4.**
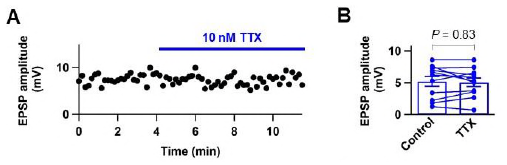
Low concentration of TTX does not affect glutamate release from presynaptic nerve terminals. (**A**) Plot of EPSP peak amplitude against time during perfusion of 10 nM TTX (blue horizontal bar). (**B**) Summary graph of the effect of 10 nM TTX on EPSP peak amplitude (control: 5.21 ± 0.78 mV, TTX: 5.06 ± 0.68, n = 11; *P* = 0.83). Data shown in **A** and example traces in **Figure 5E** were taken from the same cell. Filled circles represent data from individual experiments. Bars indicate mean ± SEM. Lines connect data points from the same experiment.

## References

1. Alme CB, Buzzetti RA, Marrone DF, Leutgeb JK, Chawla MK, Schaner MJ, Bohanick JD, Khoboko T, Leutgeb S, Moser EI, Moser MB. McNaughton BL, Barnes CA. 2010. Hippocampal granule cells opt for early retirement. Hippocampus 20:1109–1123. doi: 10.1002/hipo.20810.

2. Bliss TV, Lomo T. 1973. Long-lasting potentiation of synaptic transmission in the dentate area of the anaesthetized rabbit following stimulation of the perforant path. The Journal of Physiology 232:331–356.

3. Birnbaum SG, Varga AW, Yuan LL, Anderson AE, Sweatt JD, Schrader LA. 2004. Structure and function of Kv4-family transient potassium channels. Physiological Review 84:803–833. doi: 10.1152/physrev.00039.2003.

4. Buzsaki G. 2002. Theta oscillations in the hippocampus. Neuron 33:325–340.

5. Brunner J, Szabadics J. 2016. Analogue modulation of back-propagating action potentials enables dendritic hybrid signalling. Nature Communications 7:13033. doi: 10.1038/ncomms13033.

6. Dan Y, Poo MM. 2006. Spike timing-dependent plasticity: from synapse to perception. Physiological Reviews 86:1033–1048.

7. Diamantaki M, Frey M, Berens P, Preston-Ferrer P, Burgalossi A. 2016. Sparse activity of identified dentate granule cells during spatial exploration. Elife 5:e20252. doi: 10.7554/eLife.20252.

8. Dieni CV, Nietz AK, Panichi R, Wadiche JI, Overstreet-Wadiche L. 2013. Distinct determinants of sparse activation during granule cell maturation. The Journal of Neuroscience 33:19131–19142. doi: 10.1523/JNEUROSCI.2289–13.2013.

9. Dieni CV, Panichi R, Aimone JB, Kuo CT, Wadiche JI, Overstreet-Wadiche L. 2016. Low excitatory innervation balances high intrinsic excitability of immature dentate neurons. Nature Communications 7:11313. doi: 10.1038/ncomms11313.

10. Feldman DE. 2012. The spike-timing dependence of plasticity. Neuron 75:556–571. doi: 10.1016/j.neuron.2012.08.001.

11. Ge S, Yang CH, Hsu KS, Ming GL, Song H. 2007. A critical period for enhanced synaptic plasticity in newly generated neurons of the adult brain. Neuron 54:559–566.

12. Golding NL, Spruston N. 1998. Dendritic sodium spikes are variable triggers of axonal action potentials in hippocampal CA1 pyramidal neurons. Neuron 21:1189–1200.

13. Golding NL, Staff NP, Spruston N. 2002. Dendritic spikes as a mechanism for cooperative long-term potentiation. Nature 418:326–331.

14. Guzman SJ, Schlogl A, Schmidt-Hieber C. 2014. Stimfit: quantifying electrophysiological data with Python. Frontiers in Neuroinformatics 8:16. doi: 10.3389/fninf.2014.00016.

15. Hama K, Arii T, Kosaka T. 1989. Three-dimensional morphometrical study of dendritic spines of the granule cell in the rat dentate gyrus with HVEM stereo images. Journal of Electron Microscopy Technique 12:80–87.

16. Hamilton TJ, Wheatley BM, Sinclair DB, Bachmann M, Larkum ME, Colmers WF. 2010. Dopamine modulates synaptic plasticity in dendrites of rat and human dentate granule cells. Proceedings of the National Academy of Sciences of the USA 10:18185–18190. doi: 10.1073/pnas.1011558107.

17. Hebb DO. 1949. The Organization of Behavior (John Wiley & Sons, 1949).

18. Hoffman DA, Magee JC, Colbert CM, Johnston D. 1997. K^+^ channel regulation of signal propagation in dendrites of hippocampal pyramidal neurons. Nature 387:869–875.

19. Holmes WR. 1989. The role of dendritic diameters in maximizing the effectiveness of synaptic inputs. Brain research 478:127–137.

20. Hu H, Jonas P. 2014. A supercritical density of Na^+^ channels ensures fast signaling in GABAergic interneuron axons. Nature Neuroscience 17:686–693. doi: 10.1038/nn.3678.

21. Jarsky T, Roxin A, Kath WL, Spruston N. 2005. Conditional dendritic spike propagation following distal synaptic activation of hippocampal CA1 pyramidal neurons. Nature Neuroscience 8:1667–1676.

22. Kim S, Guzman SJ, Hu H, Jonas P. 2012. Active dendrites support efficient initiation of dendritic spikes in hippocampal CA3 pyramidal neurons. Nature Neuroscience 15:600–606. doi: 10.1038/nn.3060.

23. Kim S. 2014. Action potential modulation in CA1 pyramidal neuron axons facilitates OLM interneuron activation in recurrent inhibitory microcircuits of rat hippocampus. PLoS One 9:e113124. doi: 10.1371/journal.pone.0113124.

24. Kim Y, Hsu CL, Cembrowski MS, Mensh BD, Spruston N. 2015. Dendritic sodium spikes are required for long-term potentiation at distal synapses on hippocampal pyramidal neurons. Elife 4:e06414. doi: 10.7554/eLife.06414.

25. Krueppel R, Remy S, Beck H. 2011. Dendritic integration in hippocampal dentate granule cells. Neuron 71:512–528. doi: 10.1016/j.neuron.2011.05.043.

26. Levy WB, Steward O. 1983. Temporal contiguity requirements for long-term associative potentiation 艠 depression in the hippocampus. Neuroscience 8:791–797.

27. Letzkus JJ, Kampa BM, Stuart GJ. 2006. Learning rules for spike timing-dependent plasticity depend on dendritic synapse location. The Journal of Neuroscience 26:10420–10429. DOI: 10.1523/JNEUROSCI.2650–06.2006.

28. Lin YW, Yang HW, Wang HJ, Gong CL, Chiu TH, Min MY. 2006. Spike-timing-dependent plasticity at resting and conditioned lateral perforant path synapses on granule cells in the dentate gyrus: different roles of N-methyl-D-aspartate and group I metabotropic glutamate receptors. The European Journal of Neuroscience 23:2362–2374.

29. Lisman JE. 1999. Relating hippocampal circuitry to function: recall of memory sequences by reciprocal dentate–CA3 interactions. Neuron 22:233–242.

30. Lopez-Rojas J, Heine M, Kreutz MR. 2016. Plasticity of intrinsic excitability in mature granule cells of the dentate gyrus. Scientific Reports 6:21615 doi: 10.1038/srep21615.

31. Losonczy A, Magee JC. 2006. Integrative properties of radial oblique dendrites in hippocampal CA1 pyramidal neurons. Neuron 50:291–307.

32. Magee JC, Johnston D. 1997. A synaptically controlled, associative signal for Hebbian plasticity in hippocampal neurons. Science 275:209–213.

33. Marr D. 1971. Simple memory: a theory for archicortex. Philos. Trans. R. Soc. Lond. B Biol. Sci. 262:23–81.

34. McHugh TJ, Jones MW, Quinn JJ, Balthasar N, Coppari R, Elmquist JK, Lowell BB, Fanselow MS, Wilson MA, Tonegawa S. 2007. Dentate gyrus NMDA receptors mediate rapid pattern separation in the hippocampal network. Science 317:94–99.

35. McNaughton BL, Douglas RM, Goddard GV. 1978. Synaptic enhancement in fascia dentata: cooperativity among coactive afferents. Brain Research 157:277–293.

36. McNaughton BL, Morris RGM. 1987. Hippocampal synaptic enhancement and information storage within a distributed memory system. Trends in Neuroscience 10:408–415.

37. Menegola M, Misonou H, Vacher H, Trimmer JS. 2008. Dendritic A-type potassium channel subunit expression in CA1 hippocampal interneurons. Neuroscience. 154:953–964. doi: 10.1016/j.neuroscience.2008.04.022

38. Mishra RK, Kim S, Guzman SJ, Jonas P. 2016. Symmetric spike timing-dependent plasticity at CA3-CA3 synapses optimizes storage and recall in autoassociative networks. Nature Communications 7:11552. doi: 10.1038/ncomms11552.

39. Monaghan MM, Menegola M, Vacher H, Rhodes KJ, Trimmer JS. 2008. Altered expression and localization of hippocampal A-type potassium channel subunits in the pilocarpine-induced model of temporal lobe epilepsy. Neuroscience. 156:550–562. doi: 10.1016/j.neuroscience.2008.07.057

40. Mongiat LA, Espósito MS, Lombardi G, Schinder AF. 2009. Reliable activation of immature neurons in the adult hippocampus. PLoS One 4:e5320. doi: 10.1371/journal.pone.0005320.

41. Nevian T, Larkum ME, Polsky A, Schiller J. 2007. Properties of basal dendrites of layer 5 pyramidal neurons: a direct patch-clamp recording study. Nature Neuroscience 10:206–214. doi: 10.1038/nn1826

42. O’Neill J, Senior TJ, Allen K, Huxter JR, Csicsvari J. 2008. Reactivation of experience-dependent cell assembly patterns in the hippocampus. Nature Neuroscience 11:209–215.

43. Pernía-Andrade AJ, Jonas P. 2014. Theta-gamma-modulated synaptic currents in hippocampal granule cells *in vivo* define a mechanism for network oscillations. Neuron 81:140–152. doi: 10.1016/j.neuron.2013.09.046.

44. Ryan TJ, Roy DS, Pignatelli M, Arons A, Tonegawa S. 2015. Engram cells retain memory under retrograde amnesia. Science 348:1007–1013. doi: 10.1126/science.aaa5542.

45. Schiller J, Major G, Koester HJ, Schiller Y. 2000. NMDA spikes in the basal dendrites of cortical pyramidal neurons. Nature 404:285–289. doi: 10.1038/35005094.

46. Schmidt-Hieber C, Bischofberger J. 2010. Fast sodium channel gating supports localized and efficient axonal action potential initiation. The Journal of Neuroscience 30:10233–10242. doi: 10.1523/JNEUROSCI.6335–09.2010.

47. Schmidt-Hieber C, Jonas P, Bischofberger J. 2004. Enhanced synaptic plasticity in newly generated granule cells of the adult hippocampus. Nature 429:184–187.

48. Schmidt-Hieber C, Jonas P, Bischofberger J. 2007. Subthreshold dendritic signal processing and coincidence detection in dentate gyrus granule cells. The Journal of Neuroscience 27:8430–8441.

49. Schmidt-Hieber C, Wei H, Häusser M. 2014. Synaptic mechanisms of sparse activity in hippocampal granule cells during mouse navigation. Society for Neuroscience Abstract.

50. Scharfman HE, Schwartzkroin PA. 1990. Responses of cells of the rat fascia dentate to prolonged stimulation of the perforant path: sensitivity of hilar cells and changes in granule cell excitability. Neuroscience 35:491–504.

51. Senzai Y, Buzsáki G. 2017. Physiological Properties and Behavioral Correlates of Hippocampal Granule Cells and Mossy Cells. Neuron 93:691–704. doi: 10.1016/j.neuron.2016.12.011.

52. Sjöström PJ, Turrigiano GG, Nelson SB. 2001. Rate, timing, and cooperativity jointly determine cortical synaptic plasticity. Neuron 32:1149–1164.

53. Sjöström PJ, Rancz EA, Roth A, Häusser M. 2008. Dendritic excitability and synaptic plasticity. Physiological Reviews 88:769–840. doi: 10.1152/physrev.00016.2007.

54. Spruston N, Schiller Y, Stuart G, Sakmann B. 1995. Activity-dependent action potential invasion and calcium influx into hippocampal CA1 dendrites. Science 268:297–300.

55. Stuart GJ, Häusser M. 2001. Dendritic coincidence detection of EPSPs and action potentials. Nature Neuroscience 4:63–71.

56. Stuart GJ, Schiller J, Sakmann B. 1997. Action potential initiation and propagation in rat neocortical pyramidal neurons. The Journal of Physiology 505: 617–632.

57. Stuart GJ, Spruston N. 2015. Dendritic integration: 60 years of progress. Nature Neuroscience 18:1713–1721. doi: 10.1038/nn.4157.

58. Temprana SG, Mongiat LA, Yang SM, Trinchero MF, Alvarez DD, Kropff E, Giacomini D, Beltramone N, Lanuza GM, Schinder AF. 2015. Delayed coupling to feedback inhibition during a critical period for the integration of adult-born granule cells. Neuron 85:116–130. doi: 10.1016/j.neuron.2014.11.023.

59. Treves A, Rolls ET. 1994. Computational analysis of the role of the hippocampus in memory. Hippocampus. 4:374–391.

60. Vyleta NP, Borges-Merjane C, Jonas P. 2016. Plasticity-dependent, full detonation at hippocampal mossy fiber-CA3 pyramidal neuron synapses. Elife 5:e17977. doi: 10.7554/eLife.17977.

61. Yang K, Dani, JA. 2014. Dopamine d1 and d5 receptors modulate spike timing-dependent plasticity at medial perforant path to dentate granule cell synapses. The Journal of Neuroscience 34:15888–15897. doi: 10.1523/jneurosci.2400–14.2014.

